# Single-cell gene regulatory network reconstruction and key regulator identification using a dual-channel fusion graph convolutional network

**DOI:** 10.64898/2026.06.05.730394

**Authors:** Rongmei Tang, JianPing Liu, Pengfei Zhang, Xujun Liang

## Abstract

**Background and objective:** Gene regulatory networks are formed by complex regulatory relationships between transcription factors and their target genes. A systematic understanding of these regulatory relationships is crucial for deciphering the molecular mechanisms that underlie cell state transitions under physiological and pathological conditions. Single-cell expression data can reveal cell-type-specific transcriptional regulation, and computational methods have recently been developed to infer gene regulatory networks from single-cell transcriptomics and prior regulatory knowledge. However, existing methods could not explore the common and specific information in expression correlations and prior regulatory knowledge, which can adversely affect prediction performance.

**Methods:** We propose a novel method for inferring gene regulatory networks from single-cell RNA sequencing data. The proposed method consists of dual-channel graph neural networks and a weight-shared common graph neural network, enabling effective fusion of prior regulatory knowledge with gene co-expression patterns. Furthermore, we formulate a new computational framework built upon the proposed algorithm, which integrates differential gene expression profiles and regulatory changes to identify key regulators that distinguish different cell states.

**Results:** Experimental results demonstrate that our method significantly improves the accuracy of regulatory inference across multiple datasets, outperforming other state-of-the-art approaches. Our method also exhibits robustness to noise and missing data. Analysis of two single-cell expression datasets suggests that the proposed framework could help identify key regulators involved in tumor metastasis and drug resistance.

**Conclusion:** These results indicate that the proposed method could advance the understanding of the biological mechanisms underlying diseases by reconstructing single-cell gene regulatory networks and identifying key regulators across different cell states.

## 1. Introduction

Regulation of gene expression is crucial for cells with the same DNA sequence to perform different functions and display different morphologies. In eukaryotic organisms, gene expression can be regulated at multiple levels. An important gene expression regulation mechanism involves interactions between transcription factors (TF) and the cis-regulatory regions of target genes. The regulatory relationships between transcription factors and their targets are highly complex and play a vital role in many biological processes, such as cell differentiation and disease development[1]. Consequently, it is critical to develop a comprehensive understanding of these regulatory relationships.

The complex regulatory circuit of transcription factors and target genes in a cell could be described as a biological network, called the gene regulatory network (GRN). The edges in GRN denote the regulatory interactions between TFs and target genes. GRN offers a systemic perspective on TF-regulated gene expression. As a result, it is postulated that GRNs could be inferred from the observed gene expression profiles. There are many previous studies that attempted to predict the edges in GRNs using bulk gene expression data[2]. With rapid advances in single-cell RNA sequencing(scRNA-seq) technology, scRNA-seq data have been extensively leveraged for the reconstruction of GRN architectures. scRNA-seq data constitute a particularly promising resource for inferring GRN, owing to their distinct advantage over bulk transcriptomic data. This advantage comes from the cell-type-specific nature of TF-regulated gene expression [3]. Compared with bulk transcriptomic analyzes, scRNA-seq data more accurately capture cellular heterogeneity and dynamic changes in gene expression patterns. Consequently, scRNA-seq data facilitate the identification of cell-specific transcriptional regulatory relationships, thereby advancing the comprehension of cellular processes within distinct cell populations[4].

Recently, many computational approaches have been developed to infer the structures of GRN from scRNA-seq data. Consistent with the inference of regulatory interactions from bulk gene expression profiles, a primary strategy for deducing regulatory networks from single-cell expression data relies on gene co-expression relationships. Gene co-expression relationships refer to the correlation among multiple genes in their expression patterns[5]. The basic assumption is that co-expressed genes may be regulated by common TFs; alternatively, co-expressed TFs and target genes may engage in regulatory interactions. As a pioneering work, GENIE3 predicts the expression profile of the target gene based on all other genes using ensemble regression trees and reconstructs GRNs through feature selection[6]. GRNBoost2 is based on a concept similar to that of GENIE3, but employs gradient boosting machines to accelerate predictions for large gene expression data[7]. LEAP introduces pseudo-time to estimate the expression correlation in the scRNA-seq data, thus improving GRN inference[3]. SCENIC integrates GENIE3, which is used for co-expression analysis, with motif enrichment analysis to identify regulatory relationships between TFs and their targets[8]. Compared with linear models and tree-based methods, deep learning models excel at capturing complex nonlinear relationships within the data. Shu et al. developed a neural network-based structural equation model for GRN inference from scRNA-seq data[9]. Mao et al. proposed an unsupervised deep learning method to infer GRN through a graphical attention autoencoder[10]. Wang et al. proposed a graph autoencoder-based model named DeepRIG, which leverages weighted gene co-expression networks and global regulatory structures to infer GRNs from single-cell RNA-seq data[11].

The aforementioned methods only use single-cell gene expression data as inputs for GRN inference. Although this strategy significantly broadens the scope of application of these gene regulatory inference methods, incorporating supervised information could further improve the accuracy of the inference results. The records from the databases of the regulation of the TF-target and the ChIP-seq data provide useful supervised information. For example, DeepDRIM extracts known TF-target relationships from cell-type-specific ChIP-seq data, and uses them as supervised information to train a deep learning model for reconstructing GRNs[12]. GraphTGI is an attention-based graph embedding autoencoder model that predicts TF-target interactions by learning topological patterns within known regulatory networks[13]. In the work of Chen et al., a supervised graph attention network named GENELink was designed, which integrated scRNA-seq data and prior knowledge of gene interactions to predict gene regulatory relationships[14]. In another work, Mao et al. developed GNNLink, a supervised graph neural network framework that reconstructs GRNs from scRNA-seq data by formulating it as a link prediction task[10]. Wu et al. introduced MHHGRN, a multiview hierarchical hypergraph convolutional network designed to integrate gene expression and prior regulatory knowledge for modeling GRNs[15]. In the work of Gan et al., the global structure of regulatory relationships was captured via top-k grouping, and GraphSAGE was used to aggregate contextual information for GRN inference[16].

Although the aforementioned supervised learning methods are capable of integrating gene expression data with known gene regulatory relationships and demonstrate the potential to improve the performance of GRN inference,these studies still have some limitations. It is observed that existing methods cannot adequately integrate the gene expression data with known TF-target relationships. On the one hand, due to the dropout effect, scRNA-seq data are noisy and sparse[17], and expression correlations are insufficient evidence to support regulatory relationships. Moreover, knowledge of the TF-target relationships is also incomplete. The question of how to integrate information from the co-expression data and the prior knowledge remains a problem that needs to be resolved. Furthermore, most existing methods only propose the GRN inference model, but do not provide a framework for applying the GRN inference method to identify key regulators in diverse biological contexts.

In this study, we propose a novel GRN inference model that integrates scRNA-seq data with prior knowledge of TF-target interactions. Specifically, our approach extracts gene co-expression patterns from scRNA-seq data and employs two distinct graph convolutional networks (GCNs) to independently learn the topological structures of the co-expression graph and the prior knowledge graph. Subsequently, a weighted-shared GCN is applied, coupled with a consistency constraint, to integrate information from both sources. In this way, our framework effectively integrates complementary and biologically consistent information derived from scRNA-seq data and established TF-target relationships. To evaluate the proposed method, we conduct comprehensive comparisons with 11 state-of-the-art GRN inference algorithms. Benchmarking experiments demonstrate that our approach achieves superior performance across multiple datasets. The proposed method also exhibits strong robustness (e.g., against noise and missing data). Building upon this GRN inference algorithm, we further develop a computational framework to identify key regulatory factors governing cell state transitions, which integrates both differential regulatory interactions and gene expression changes, enabling systematic identification of important regulators that drive phenotypic alterations. To validate our framework, we applied it to two scRNA-seq datasets. The analysis revealed novel regulatory mechanisms and context-specific master regulators, confirming the framework’s utility for extracting actionable biological knowledge from single-cell data.

## 2. Materials and methods

### 2.1. Data preparation and processing

This work uses the BEELINE benchmark to evaluate the performance of the GRN inference algorithms [18]. The dataset comprises two parts. The first part covers the experimental single-cell RNA sequencing data. The scRNA-seq data consist of seven cell types. Two human cell lines are human embryonic stem cells (hESC)[19] and human mature hepatocytes (hHEP) [20]. Four cell lines from mice are mouse embryonic stem cells (mESC)[21], mouse dendritic cells (mDC)[22], mouse hematopoietic stem cells with erythroid lineage (mHSC-E), mouse hematopoietic stem cells with macrophage-granulocyte lineage (mHSC-GM), and mouse hematopoietic stem cells with lymphoid lineage (mHSC-L) [23]. These RNA-seq data originate from Chen’s work [14], and are preprocessed using the same pipeline as the BEELINE framework [18]. Specifically, the 500 and 1000 most significantly varying genes and TFs, with a corrected p-value cut-off of 0.01, are retained for GRN reconstruction. The second part of the datasets comprises ground-truth networks. The functional interactions between genes are extracted from the STRING database[24]. The nonspecific TF-target networks are derived by integrating records from multiple databases sources, including DoRothEA[25], RegNetwork [26] and TRRUST[27]. Cell-type-specific gene regulatory networks are constructed from ChIP-seq data for special cell types. Additionally, a network generated through Loss/Gain of Function (LOF/GOF) experiments is available for mESC. Table S1 presents the statistics of the scRNA-seq datasets and the ground-truth networks.

### 2.2. Proposed method

In this work, the GRN inference task is modeled as a link prediction problem. To effectively fuse gene expression information from scRNA-seq data and the topological structure informed by known TF-target relationships, we propose a novel neural network model named dual-channel fusion graph neural network (DFGNN). Our model consists of two parts. The first part comprises two graph convolutional networks to separately mine the coexpression patterns in scRNA-seq data and the topological structures in the known TF-target network. Subsequently, the node representations from these two channels are fed into the second part of the model, which is also a graph convolutional network, to learn the consistency between gene expression and prior knowledge. After that, the specific representations derived from the first part and the common representations from the second part are fused to predict the missing links between the TFs and the target genes. Figure 1 shows the framework of the proposed model, and the detailed structure of the model is described in the following.

**Figure 1:**
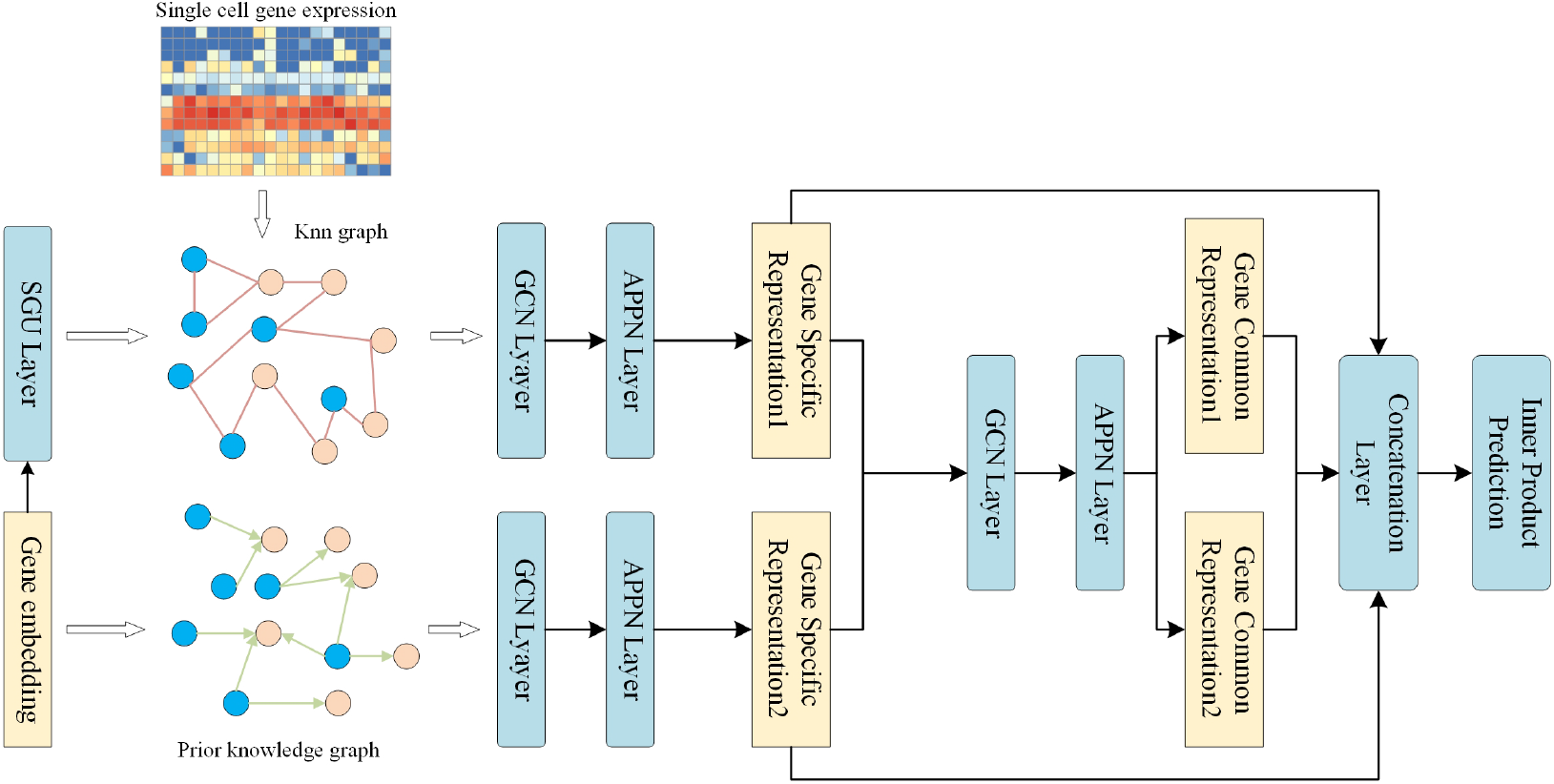
The overview of the proposed model.

#### 2.2.1. Construction of dual channel graph neural network

After preprocessing the scRNA-seq data as described in 2.1, we obtain the gene expression data *X* ∈ ℝ^*n×m*^, where *n* is the number of genes and *m* is the number of cells. To explore the co-expression patterns of genes, we calculate the cosine similarity matrix *S* ∈ ℝ^*n×n*^ of gene expression. The element *S*_*ij*_ in *S* is:

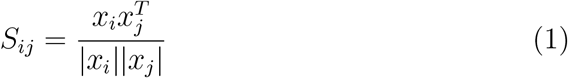

where *x*_*i*_ and *x*_*j*_ are the *i*-th row and *j*-th row of *X*, and |*x*| denotes the norm of *x*. Then the graph of the k-nearest neighbors (kNN) is constructed as *G*_*e*_, and the adjacency matrix *A*_*e*_ is:

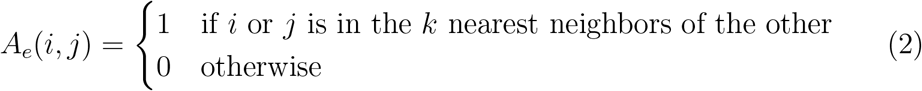

where the nearest neighbors are derived from the similarity matrix *S*.

Next, we use the randomly initialized embedding matrix *X*_*e*_ ∈ ℝ^*n×d*^ as the node features, where *d* denotes the dimension of the node features. With *X* and *A*_*e*_ as input, a two-layer graph convolutional module is designed to capture the structure information in the feature space of gene expression. The first layer can be expressed as:

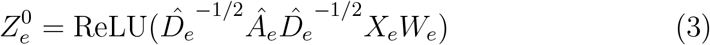

where ReLU denotes Rectified Linear Unit, *W*_*e*_ is a learnable weight matrix, Â_*e*_ = *A*_*e*_ + *I*, and *I* is an identity matrix. 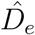 is the diagonal degree matrix, and 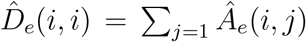. The second layer is an approximate personalized propagation of neural prediction layer[28]. It is formulated as follows:

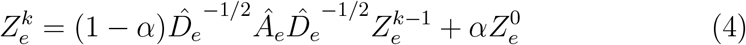

where *α* ∈ (0, 1] is the probability of restart. This layer can be iterated *K* times. *α* and *K* are hyperparameters. 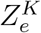 represents the node representa-tions learned from the gene expression graph *G*_*e*_. Thus, as shown in equation (3), the first layer of the graph convolution network transforms the embedding of nodes and aggregates the local information of the graph. The second layer (as shown in equation (4)) of personalized propagation can explore a larger neighborhood in the graph and avoid over-smoothing.

The known TF-target relationships constitute the second graph *G*_*p*_, and its adjacency matrix is denoted as *A*_*p*_. Since the information in the prior knowledge graph *G*_*p*_ could differ from that in the gene expression graph *G*_*e*_, we use self-gating units to adapt the embedding matrix *X*_*e*_:

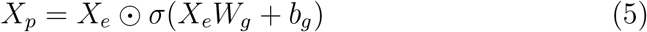

where *W*_*g*_ ∈ ℝ^*d×d*^, *b*_*g*_ ∈ ℝ^*d*^ are learnable parameters and ⊙ denotes the Hadamard product. *σ* is the sigmoid function. A two-layer graph convolution module is constructed with the same structure as described in equations (3) and (4) but with independent parameters. It takes the embedding matrix *X*_*p*_ as input, and this network module learns the representations of the nodes 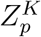 based on the graph *G*_*p*_:

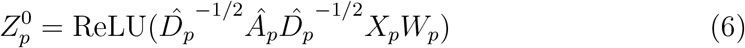

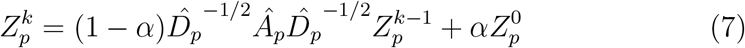

where *W*_*p*_ is a learnable weight matrix. *Â*_*p*_ = *A*_*p*_ + *I, I* is an identity matrix. 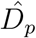 is the diagonal degree matrix and 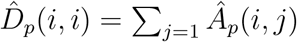.

#### 2.2.2. Construction of fusion graph neural network

The dual-channel graph neural network described above extracts information from gene co-expression patterns and prior knowledge, respectively. Subsequently, to integrate the information derived from both sources, we construct a fusion neural network inspired by the previous study[29]. The architecture of the fusion neural network is identical to that of the dual-channel graph neural network. It takes the output of the dual-channel graph neural network 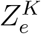 and 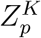 as input:

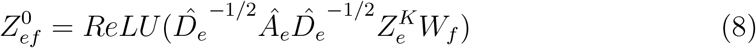

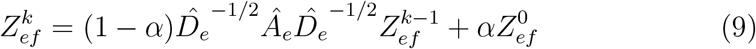

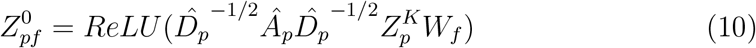

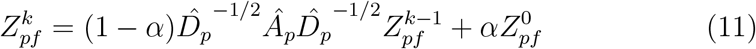

where *W*_*f*_ is a learnable weight matrix. 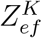 and 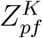 are the learned node representations. A constraint is proposed to further enhance the consistency between them:

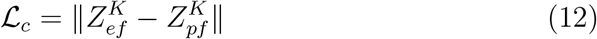

#### 2.2.3. Prediction of regulatory relationships and objective functions

After obtaining the outputs 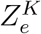 and 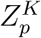 of the dual-channel graph neural network, and the outputs 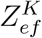 and 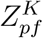 of the fusion graph neural network, we form the final node representations as:

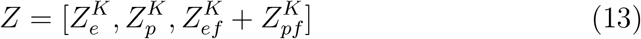

where [·] is the concatenation operation. The link between a transcription factor and a target gene is predicted by the dot product of their representations

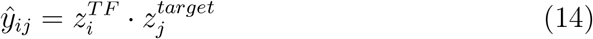

where 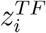 is the representation of the transcription factor *i* (i.e., the *i*-th row of the matrix *Z*), and 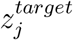 is the representation of the target gene *j* (i.e., the *j*-th row of the matrix *Z*). The known TF-target relationships *y*_*ij*_ are used as supervised information to train our model, and the supervised loss function is:

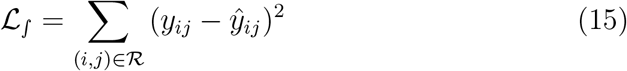

where ℛ consists of a positive and a negative link set for model training. The overall loss function is as follows:

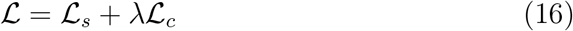

where *λ* is a hyperparameter. We utilize the Adam optimizer to update the parameters of the proposed model.

### 2.3. Data processing and key regulator detection framework for case-control comparative analysis

Building upon our GRN inference algorithm, we develop a computational framework to identify key differential regulators and regulatory effects between case and control conditions. This framework integrates both the topological differences in the GRNs and the differential gene expression profiles across case-control states. We applied this framework to two cRNA-seq datasets. The first dataset is derived from a study of human breast cancer metastasis (BCM)[30]. The GRNs of the primary tumor cells and the lung metastatic cells are constructed and compared. The second dataset is obtained from a study of multiple myeloma (MM)[31]. We compare GRNs in malignant plasma cells (PCs) between non-responder and responder patients in the combined-regimen group. The R package ‘Seurat’ is employed to preprocess these datasets[32]. For quality control of the BCM dataset, cells with fewer than 2,500 unique features or mitochondrial gene content exceeding 50% are removed. In the MM dataset, cells with fewer than 300 unique features or mitochondrial gene content exceeding 50% are filtered out. Non-plasma cells (non-PCs) in the MM dataset are excluded according to the original study. The 2,000 most variable genes are selected for each datasets, and differential expression (DE) analysis is performed using the ‘FindMarkers’ function in Seurat. We extract prior regulatory relationships from the Regnetwork database to serve as the background network[33]. Then, the gene regulatory network is constructed with the proposed method separately for primary tumor cells and lung metastatic cells in the BCM dataset, as well as for PCs from the non-responder group and the responder group in the MM dataset. To identify key regulatory factors that distinguish the control state from the case state, we defined an influence score *R*_*i*_ for each regulator *i*.

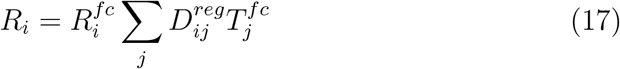

where 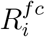 represents the logarithmic transformation of the absolute fold change value for regulator *i*, 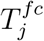 is the logarithmic transformation of the absolute fold change value for target *j*, and 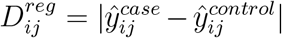 quantifies the absolute difference in interaction scores between the case and control states. The influence score *R*_*j*_ thus captures both regulatory and expression changes. Based on this score, we could prioritize all regulators according to their importance in distinguishing the control from the condition of the case. We employ GSEApy[34] to perform a pathway-enrichment analysis for top-ranked regulators.

## 3. Results

### 3.1. Experimental setting

In this work, the known relationships between pairs of transcription factors (TF) and target genes are taken as positive samples, while TF-gene pairs that are not included in the ground truth networks are taken as negative samples. The known TF-gene pairs account for only a small fraction of all possible TF-gene pairs (Table S1). Thus, the number of negative samples is much larger than the number of positive samples. In addition, the characteristic of the cell-type-specific ground truth network, which has a limited number of TFs and a higher network density, is substantially different from the other three types of ground truth networks.

Based on these observations, we adopt the data division strategy employed in the previous work[14] to evaluate the performance of the proposed model and the methods compared. Specifically, for the STRING, the non-specific, and the Loss/Gain of Function (LOF/GOF) ground-truth networks, we divide the known TF-gene pairs into three roughly equal-sized parts. One part of the samples is taken as the positive test set. We further select one-fifth of the samples from the remaining portion as the positive validation set, while the remaining four-fifths of them are designated as the positive training set. To address the data-imbalance problem, where negative samples dominate positive ones, we prioritize hard negative samples to enhance model training. For each positive TF-gene pair in the training set, we select a target gene that has no known relationship with the TF to form a new TF-gene pair as the hard negative sample for model training. The negative samples in the validation set are selected in the same way. We randomly select the negative samples for the test set from the remaining unknown relationships and ensure that the positive-to-negative ratio in the test set aligns with that of the source network.

For cell-type-specific networks, the positive target genes of each TF are divided into three roughly equal-sized groups. Two groups are used as the positive training set. 10% of the remaining samples are designated as the positive validation set, and the rest are used as the positive test set. The negative target genes of each TF are divided to form the negative training set, validation set, and test set in the same manner.

To evaluate the efficacy of the proposed method in GRN reconstruction, we compare it with several baseline methods and leading models widely used for GRN inference. As our method is based on graph neural networks, we also include some general graph learning models for comparison to demonstrate the advantage of the proposed model.

- PCC (Pearson correlation coefficient) directly computes the expression correlation between TFs and target genes. It is a simple and intuitive method for GRN inference.
- GENIE3[6] is a classic GRN inference algorithm based on regression trees.
- GRNBoost[7] uses gradient boosting machines and gene expression data to reconstruct the GRN.
- DeepWalk[35] is a random-walk-based method and serves as an efficient and general algorithm for learning node representations in a graph. We used the inner products of the node representations generated by DeepWalk to predict the links between TFs and target genes.
- mvgrl[36] is a self-supervised approach that learns node-level representations by maximizing mutual information between representations encoded from different views of the same node. We select it as the compared method because it represents another way to integrate gene co-expression and prior knowledge.
- DeepSEM[9] is a deep generative model for GRN inference, grounded in structural equation modeling.
- DeepRIG[11] utilizes a graph autoencoder model to infer TF-target regulatory relationships.
- GraphTGI[13] is a deep model composed of a graph attention-based encoder and a bilinear decoder to predict TF-target interactions.
- GENELink[14] uses a graph attention network to model the inference of TF-target interactions as a link prediction problem.
- GNNLink[37] is similar to GENELink, but uses a graph convolutional network as the backbone of its model.
- IGEGRNS[16] is a GraphSAGE-based supervised deep learning model designed to infer GRNs from scRNA-seq data.

All models are trained using the training set. The hyperparameters of our model are optimized using the validation set. The hyper-parameters of the compared methods are set as suggested in the original papers or optimized on the validation set if better performance could be obtained. The detailed settings of the hyperparameters could be found in the supplementary material. The performance of the models is evaluated on the test set. As in previous work[14, 37], we select the receiver operating characteristic curve (AUROC) and the area under the precision-recall curve (AUPRC) as metrics to evaluate the GRN inference performance of each model. To obtain a robust comparison result, we repeat the data partitioning ten times. Each partition has different samples in the training, validation, and test sets. The average performance metrics are compared.

### 3.2. Performance comparison

To evaluate the performance of the proposed model, we compare the gene regulatory network (GRN) inference capabilities of our model with those 11 other methods described above, using the 44 datasets from the BEELINE benchmark. The AUROC and AUPRC metrics of our model and the comparative methods are presented in Figure 2 (TF+500 dataset) and Figure 3 (TF+1000 dataset). To obtain more robust results, we repeated performance evaluation experiments ten times with different data divisions and performed rank-sum tests to compare the metrics of our model with those of other other methods on each data set. The statistical significance counts and the average performance improvement are shown in Table 1. It is evident that our model consistently shows better or comparable performance across the majority of datasets. For AUROC, our model significantly outperforms the second-best method, GNNLink, on 38 datasets, while is inferior to GNNLink on a single dataset (the TF+1000 STRING mHSC-L dataset, p-value*<*0.05). The average AUROC improvement over GNNLink is 3.2%. Our method achieves the highest AUROC of 0.91 on the TFs+1000 non-specific hESC and hHEP dataset, which represents a 10% improvement over GNNLink. The largest improvement in AUROC is 41.7%, which is achieved in comparison with DeepRIG. Our method demonstrates significantly superior AUROC performance compared to two classic algorithms, GRNBoost and GENIE3, with relative improvements of 37% and 36%, respectively.For AUPR, our model demonstrates significantly superior performance compared to the second-best method, GNNLink, on 34 datasets, and performs slightly worse on only 2 datasets (p-value < 0.05). The average AUPR improvement relative to GNNLink is 5.8%. Our method exhibits the largest AUPR im-provement over GNNLink in the TF+500 and TF+1000 LOF/GOF mESC datasets, reaching 25.5%. Our model significantly outperforms IGEGRNS and mvgrl on most datasets, with average AUPR improvements of 18.0% and 19.2%, respectively. Except for these three methods, the AUPR metrics of the other methods are significantly inferior to those of our model across all 44 datasets (p-value < 0.05). These findings indicate that our model yields consistently superior performance in inferring gene regulatory networks.

**Table 1:** Statistical significance counts across multiple datasets for different methods.

|  | Our model |  |  |  |  |  |  |  |
| --- | --- | --- | --- | --- | --- | --- | --- | --- |
|  | AUROC |  |  |  | AUPR |  |  |  |
| | Better | Worse | No diff. | ΔAvg | Better | Worse | No diff. | ΔAvg |
| IGEGRNS | 43 | 0 | 1 | 10.2% | 39 | 2 | 3 | 18.0% |
| GNNLink | 38 | 1 | 5 | 3.2% | 34 | 2 | 8 | 5.8% |
| GENELink | 44 | 0 | 0 | 10.1% | 44 | 0 | 0 | 15.5% |
| GraphTGI | 43 | 0 | 1 | 16.9% | 44 | 0 | 0 | 21.7% |
| DeepRIG | 44 | 0 | 0 | 41.7% | 44 | 0 | 0 | 36.1% |
| DeepSEM | 44 | 0 | 0 | 36.4% | 44 | 0 | 0 | 34.2% |
| mvgrl | 44 | 0 | 0 | 19.6% | 42 | 1 | 1 | 19.2% |
| DeepWalk | 44 | 0 | 0 | 12.8% | 44 | 0 | 0 | 24.6% |
| GRNBoost | 44 | 0 | 0 | 37.3% | 44 | 0 | 0 | 33.5% |
| GENIE3 | 44 | 0 | 0 | 36.2% | 44 | 0 | 0 | 32.6% |
| PCC | 44 | 0 | 0 | 33.0% | 44 | 0 | 0 | 30.9% |

'Better' indicates that our model's metric is significantly higher than that of the compared method. 'Worse' suggests that our model's metric is significantly lower. 'No diff.' signifies that there is no statistically significant difference. The values in the table represent the counts of such statistical differences in model performance across multiple datasets. Statistical analyses were performed using the rank-sum test; differences were considered significant at $p < 0.05$ . $\Delta$ Avg is the average performance improvement of our model compared to the comparative method.

**Figure 2:**
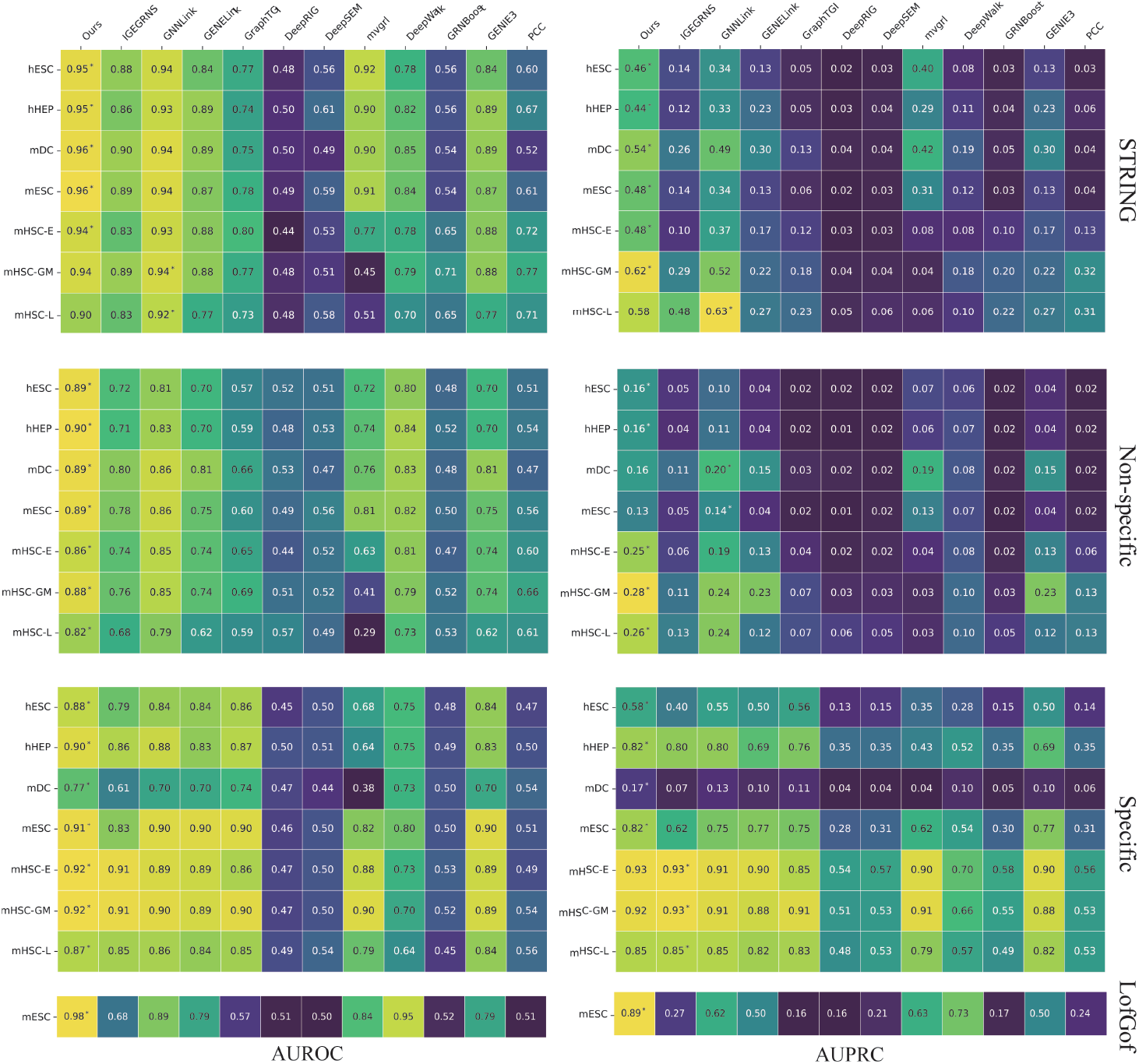
The comparison of AUROC and AUPRC for 12 methods on the TF+500 benchmark dataset. Results are averaged across 10 independent runs, and a single asterisk (*) denotes the best performance for each dataset.

**Figure 3:**
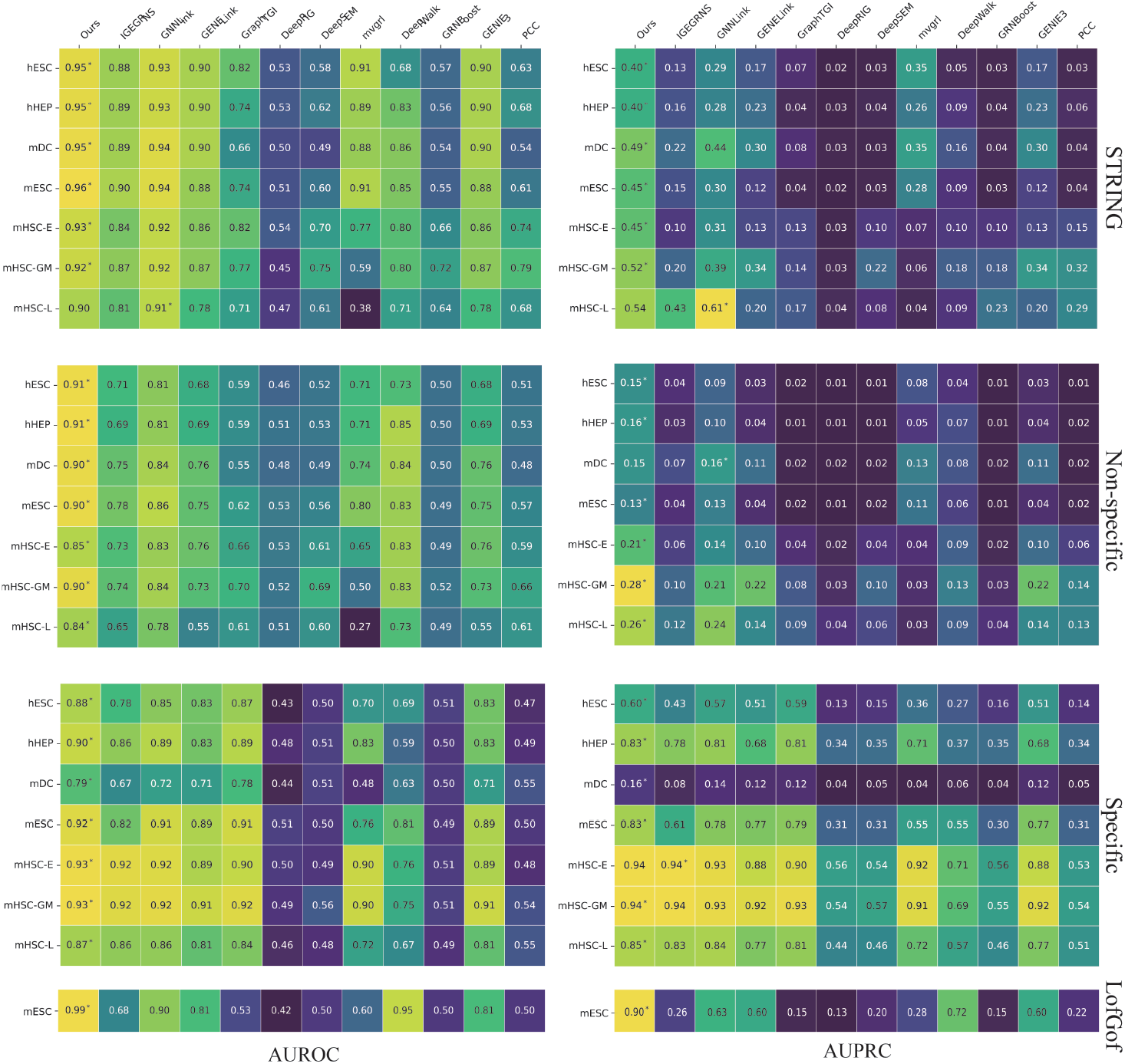
The comparison of AUROC and AUPRC for 12 methods on the TF+1000 benchmark dataset. Results are averaged across 10 independent runs, and a single asterisk (*) denotes the best performance for each dataset.

### 3.3. Sensitivity analysis of hyperparameters, data perturbation and model architecture

The performance of the proposed model can be impacted by the hyperparameters. Here, we examine the influence of several key hyperparameters. We first investigate how the learning rate and weight decay influence the prediction performance. AUROC values are compared under different settings of hyperparameters using the TF+500 non-specific and specific datasets. As shown in Figures 4A and B, the AUROC values decrease when the learning rate is too low or too high, but exhibit minimal changes when the learning rate is within the range of 1e-2 to 1e-4 across most cell types. As shown in Figures 4C and D, the weight decay affects the performance of the proposed model more obviously. The model achieves its best performance with different weight decay values for the non-specific dataset (1e-2) and the specific datasets (1e-3). The hyper-parameter *λ* in equation (16) regulates the trade-off between the two components of the final loss function. As shown in Figures 4E and F, for most datasets, our model is relatively stable when the value of *λ* is within the range of 0.5 to 5. For the non-specific TF+500 mHSC-L dataset, the AUROC values first decrease and then increase with increasing *λ*. The performance of our model is not very sensitive to the embedding dimension of node features on most datasets, as shown in Figures S1A and B.

**Figure 4:**
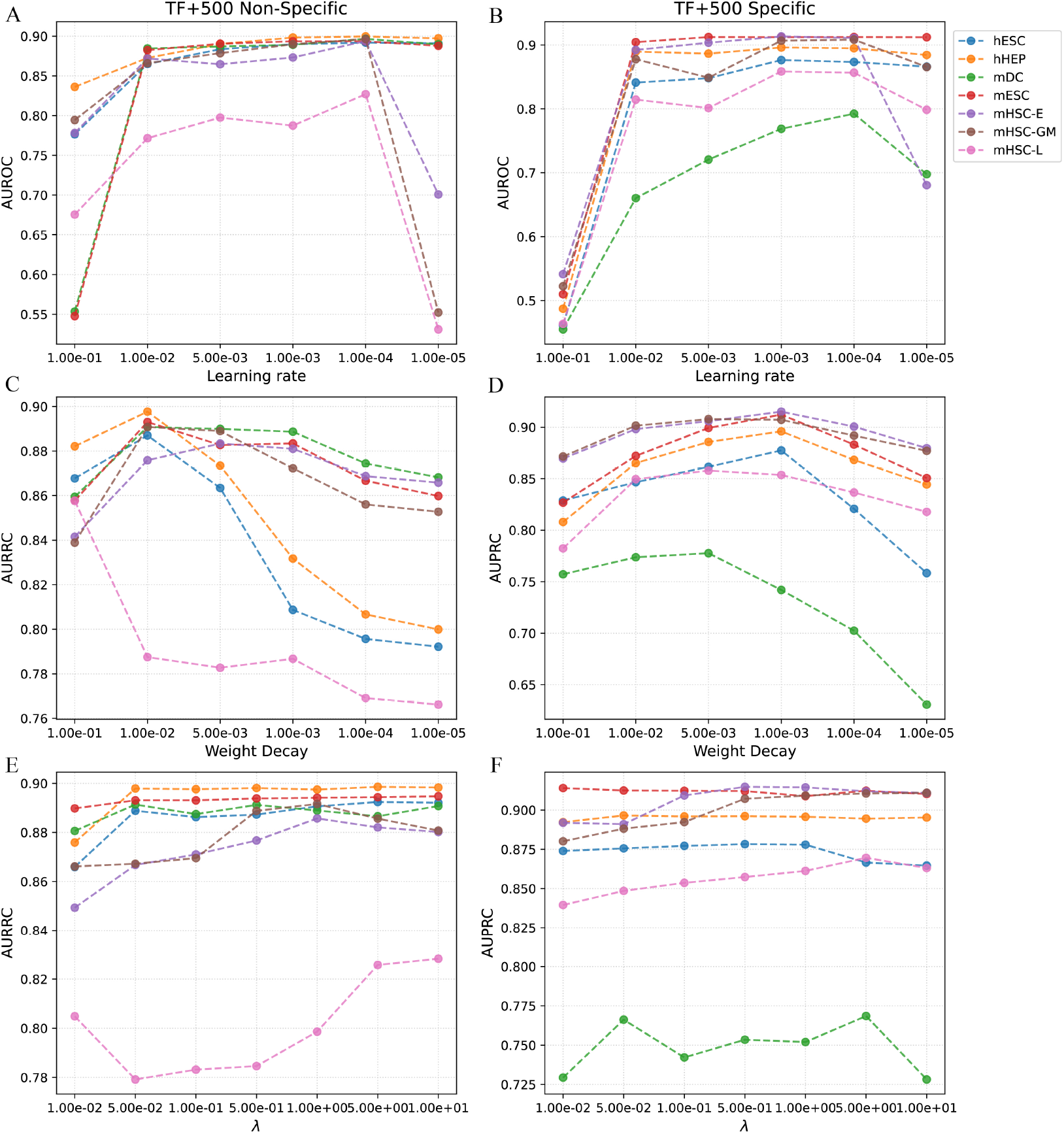
The AUROC and AUPRC values of the proposed method under different hyperparameter settings on the TF+500 non-specific and specific datasets. (A) and (B) show the influence of learning rate. (C) and (D) show the influence of weight decay. (E) and (F) show the influence of *λ*.

To test the robustness of our model, we performed data perturbation and examined the performance changes. The TF+500 specific datasets are used for these experiments. Here, we performed two types of data perturbation. The first type of perturbation is desinged to mask a portion of the known TF-target relationships. Specifically, the elements in the adjacency matrix *A*^*p*^ are randomly set to 0, and the corresponding positive samples in the training target {*y*_*ij*_}_*pos*_ are also set to 0. As shown in Figure 5A, the AUROC scores decrease as the percentage of the masked TF-target edges increases. However, when the percentage of remaining TF-target relationships exceeds 70%, the variation in AUROC scores is not greater than 0.05. It is implied that our model could still perform well when part of the known TF-target relationships is missing. The second type of data perturbation involves randomly dropping some cells in the expression data. As shown in Figure 5B, even with only half of the cells in the original expression data, our model still performs well for GRN inference. This suggests that our model could reconstruct GRNs for rare cell subpopulations with limited cell number.

**Figure 5:**
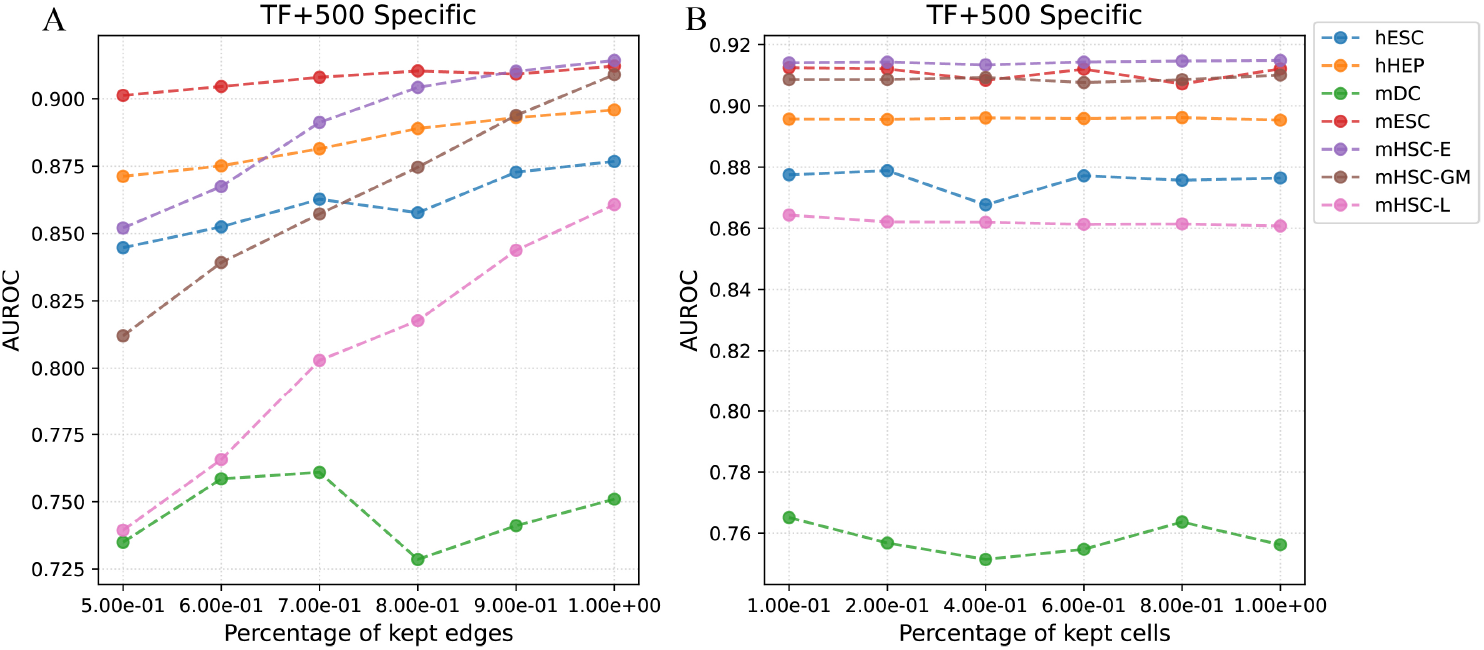
The AUROC and AUPRC values of the proposed method under different data perturbations on the TF+500 specific datasets. (A) shows the influence of masking known TF-target relationships. (B) shows the influence of cell dropping.

Next, we examine the impact of different model architectures on prediction performance. To investigate whether the consistency constraint is meaningful, we remove the constraints ℒ_*c*_ from the objective function. To investigate the effect of the approximate personalized propagation of neural predictions (APPNP), we replace it with a graph convolution layer that matches the first layer. As shown in Figure 6 and Table 2, the model without the consistency constraint performs inferior to the original model on most datasets. It also demonstrates lower performance than the model without the APPNP layer on most datasets. This confirms the effectiveness of the consistency constraint. Regarding the model without the APPNP layer, although the original model’s performance improvement is not particularly pronounced in terms of AUPRC, it still attains a superior AUROC metric on most datasets. Additionally, a comparison between Figure 6 and Figures 2 and 3 reveals that these model variants maintain competitive performance relative to the baseline method, suggesting that the basic architecture of our proposed model possesses distinct advantages.

**Table 2:** Statistical difference counts in model variants comparisons.

|  | Original model |  |  |  |  |  |  |  |
| --- | --- | --- | --- | --- | --- | --- | --- | --- |
|  | AUROC |  |  |  | AUPR |  |  |  |
| | Better | Worse | No diff. | ΔAvg | Better | Worse | No diff. | ΔAvg |
| Without APPNP | 21 | 2 | 21 | 1.0% | 10 | 10 | 24 | 0.4% |
| Without $\mathcal{L}_c$ | 31 | 1 | 12 | 2.3% | 31 | 4 | 9 | 4.3% |

**Figure 6:**
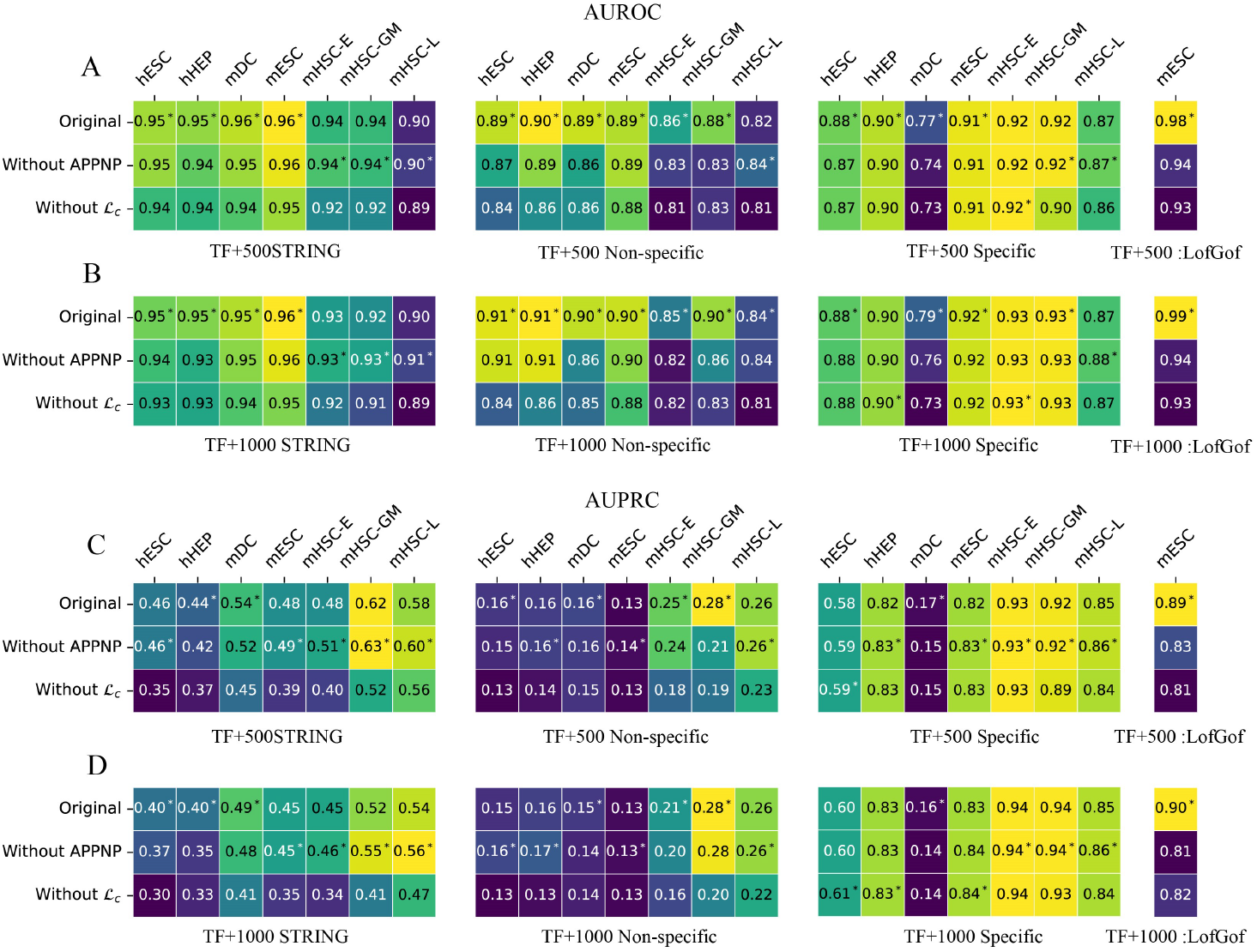
The comparison of AUROC and AUPRC across model variants on the TF+500 and TF+1000 benchmark datasets. Results are averaged over 10 independent runs, and a single asterisk (*) denotes the best performance for each dataset.

### 3.4. Application example: identification of key regulatory factor based on our GRN inference method

The differential activity of regulatory factors serves as a crucial determinant in driving cellular state transitions and maintaining homeostasis. A primary objective of reconstructing the gene regulatory network is to quantitatively identify regulators with condition-specific activities. Here, we apply our proposed method to infer GRNs that govern different cellular states. In addition, we develop a computational framework that integrates topological differences in GRNs with gene expression dynamics to estimate changes in regulator activity. As a proof of concept, we applied the proposed framework to two scRNA-seq datasets.

The first dataset consists of three patient-derived xenograft (PDX) models of triple-negative breast cancer (TNBC). Using our model, we first construct gene regulatory networks (GRNs) separately for primary tumor cells and lung metastatic cells. Then we calculate the differential expression of genes between these two groups of cells and rank the regulators according to the influence score defined in equation (17). The complete list of regulators and influence scores can be found in Supplemental Table S2 and the distribution of the influence scores is shown in Supplemental Figure S2. To investigate the biological pathways in which the most influential regulatory factors are involved, we performed a KEGG pathway enrichment analysis on the top 40 regulators ranked by their influence scores. Significantly enriched KEGG pathways include ESR-mediated signaling, cell cycle-related pathways, and Notch signaling (Figure 7A). In the metastatic progression of triple-negative breast cancer (TNBC), the Notch signaling pathway plays a central driving role and is often considered a hallmark of this subtype. It critically regulates tumor cell invasion and distant colonization[38]. Next, to demonstrate the details of differential regulation between primary tumor cells and lung metastatic cells, we selected the 100 edges with the largest changes in regulatory scores, together with their connected nodes, and constructed the differential regulatory network shown in Figure 7B. In this network, the regulatory relationship between FOS and JUN, previously established in training datasets, is found to be strengthened in lung metastatic cells according to our analysis. Previous studies have demonstrated that FOS and JUN form the AP-1 heterodimer, which plays a critical role in the invasion of breast cancer [39]. Our GRN construction algorithm predicts a novel regulatory interaction between MYC and IRF1 that is absent from the training data, and further reveals dynamic changes in this regulatory relationship. A previous study identified that in the TNBC model, MYC suppresses the expression of multiple interferon-stimulated genes, including IRF1, and promotes tumor immune escape[40].

**Figure 7:**
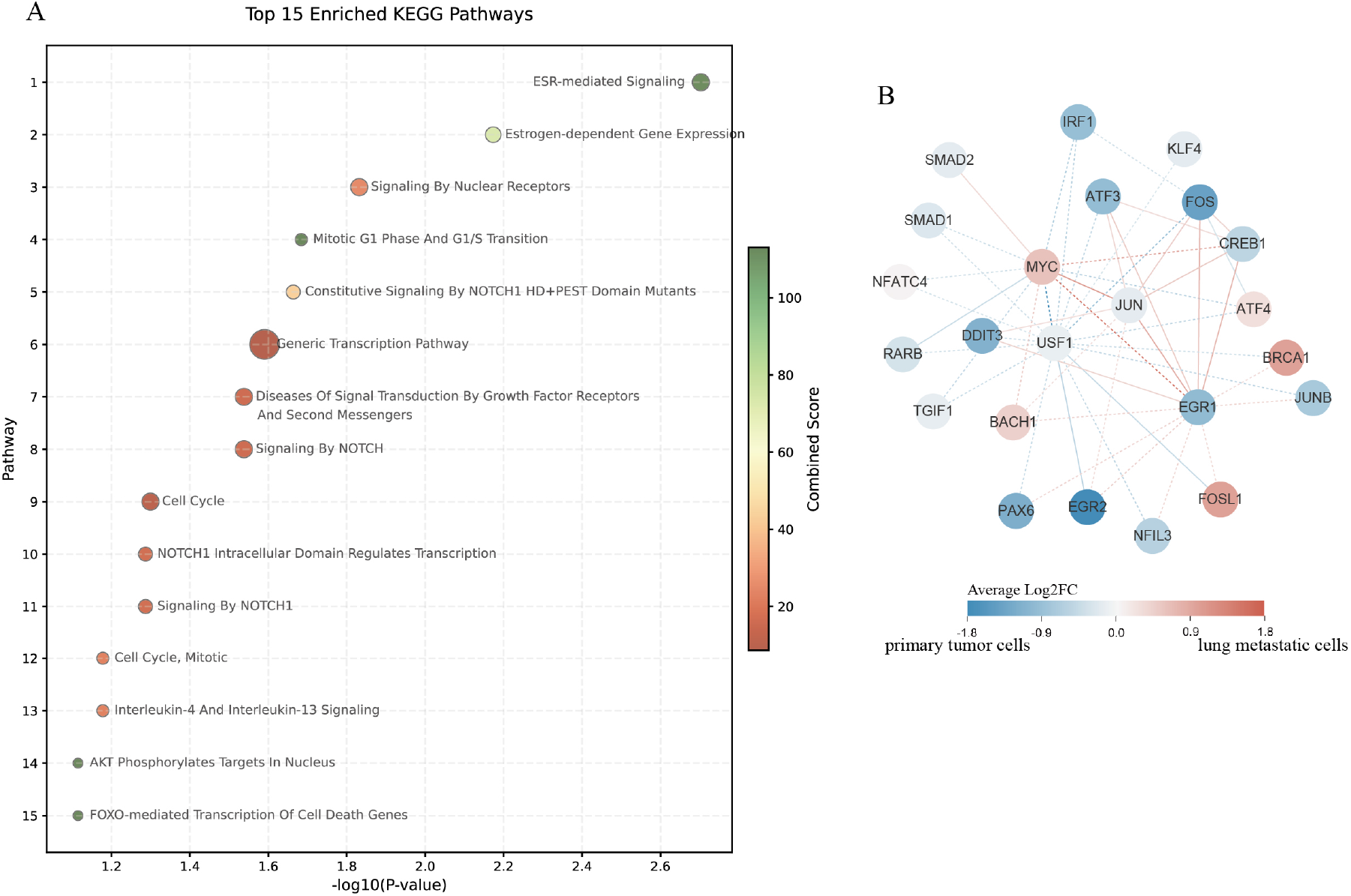
Differential regulator analysis of primary tumor cells and lung metastatic cells. (A) KEGG pathway enrichment analysis of the top-ranked regulators. The size of each dot corresponds to the number of genes linked to the respective pathway. (B) The differential regulatory network consists of the 100 edges exhibiting the most significant regulatory changes. Edge colors reflect the magnitude of regulatory score changes, whereas node colors indicate differential gene expression. Solid lines represent known interactions derived from the training data, and dashed lines denote novel predicted relationships.

The second dataset is analyzed using our method to examine the differential regulatory networks between drug-non-responding and drug-responding malignant plasma cells. The influence scores of all regulators are provided in Supplemental Table S3. Based on the distribution of influence scores (Figure S3), we performed KEGG pathway enrichment analysis on the top 50 regulators.The first 15 enriched pathways are shown in Figure 8A. Among these pathways, cell cycle-related pathways are not only a core driver of tumor proliferation but also a critical mechanism underlying drug resistance in multiple myeloma[41]. The Wnt signaling pathway is closely associated with drug resistance in multiple myeloma[42]. This connection manifests itself not only through intrinsic mechanisms within myeloma cells but also through their interactions with the bone marrow microenvironment[43]. The differential regulatory network, comprising 100 edges with the most significant changes in regulatory scores and their connected nodes, is shown in Figure 8B. In multiple myeloma, the reciprocal regulation between MAX and MYC serves as one of the main mechanisms that drive disease progression and confer drug resistance[44]. Our method predicts a novel association between MAX and FOXO3. A previous study showed that FOXO3 is a critical regulator of drug resistance in multiple myeloma[45]. MAX could directly bind to the FOXO3 promoter region to modulate its transcription[46]. The examples above demonstrate that the GRN inference algorithm proposed in this study, along with its derived differential regulatory network identification method, successfully uncovers condition-specific cellular regulatory networks and pinpoints key regulatory factors.

**Figure 8:**
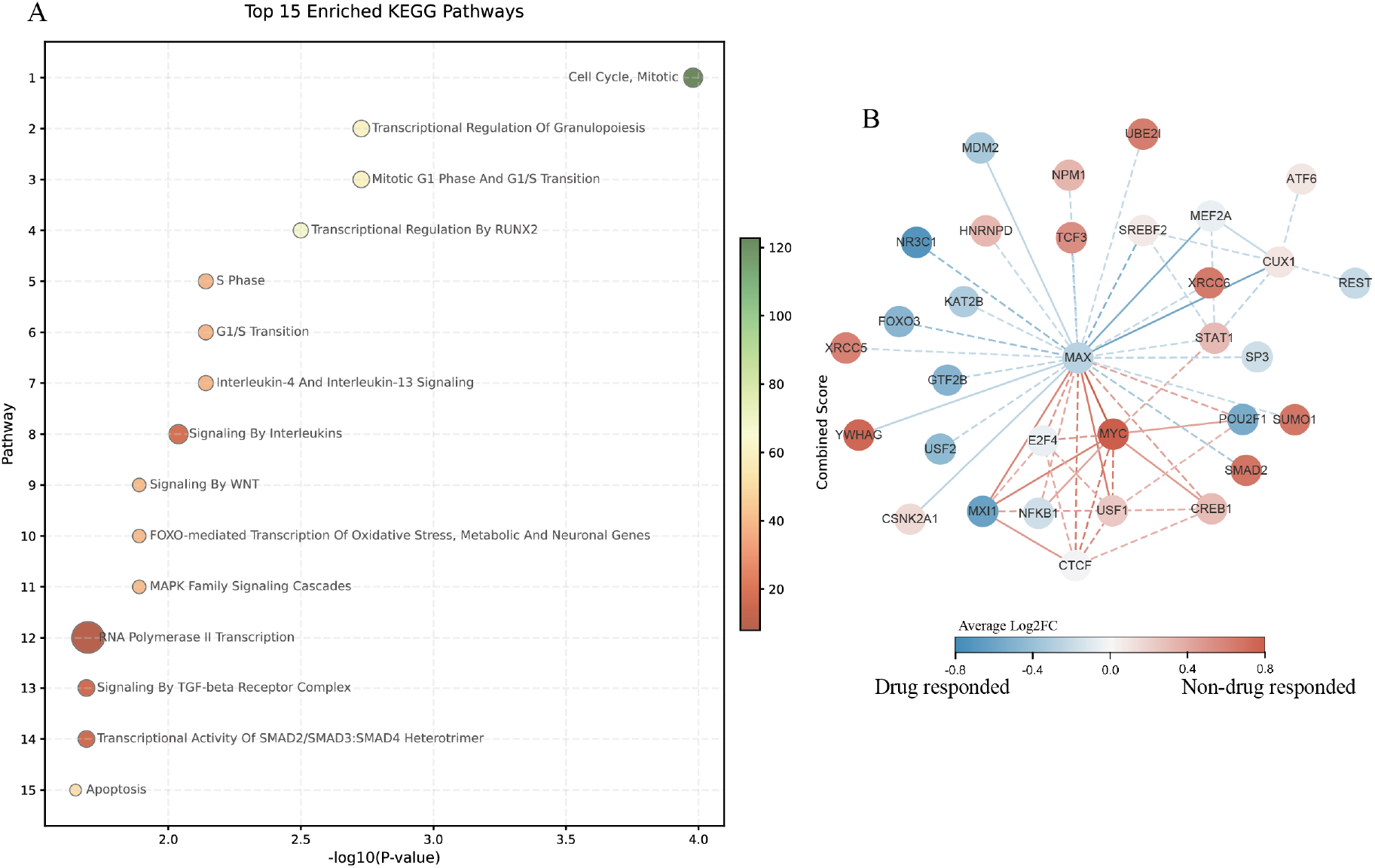
Differential regulator analysis of drug-non-responded and responded malignant plasma cells. (A) KEGG pathway enrichment analysis of the top-ranked regulators. Dot size corresponds to the number of genes linked to the respective pathway. (B) The differential regulatory network consists of the 100 edges exhibiting the most significant regulatory changes. Edge colors reflect the magnitude of regulatory score changes, whereas node colors indicate differential gene expression. Solid lines represent known interactions derived from the training data, and dashed lines denote novel predicted relationships.

## 4. Discussion

Gene regulatory networks (GRNs) are complex network systems composed of interactions among target genes, transcription factors, and regulatory elements (e.g., promoters, enhancers). Inferring GRNs from observed data (e.g., gene expression profiles) represents a critical approach to elucidate biological processes and uncover the mechanisms underlying phenotypic outcomes. Single cell sequencing data facilitate more precise and cell-type-specific inference of GRNs. Moreover, with prior knowledge of TF-target relationships derived from databases or ChIP-Seq data, the GRN inference problem could be modeled as a link prediction task using supervised learning algorithms. Despite recent advances, the integration of scRNA-seq data with prior regulatory networks remains a critical unmet challenge. In addition, inferred networks should be applied to discover key regulatory factors, offering actionable insights for subsequent experimental research.

In this study, we propose a novel algorithm for inferring GRNs by integrating scRNA-seq data with prior knowledge. Unlike previous approaches that use gene expression profiles as node features and predefined regulatory relationships as topological structures, our method adopts a multi-channel neural network architecture, inspired by the previous study[29] . We design a network module composed of the conventional graph convolutional layer and the personalized propagation layer to capture local information and then extend to a larger neighborhood. We use two specific network modules and one common network module to integrate co-expression patterns with prior topological network. This model enables the performance improvement of GRN inference compared with several state-of-the-art methods. The algorithm maintains robustness with partial prior knowledge or missing expression data.

Complex phenotypical transitions, such as tumor metastasis or drug resistance, often involve multi-gene and multi-layer cooperative changes. In this study, we introduce a novel computational framework that relies on our GRN inference algorithm to jointly analyze changes in regulatory structure and differential gene expression. The experimental results show that this framework can characterize distinct disease states (e.g., metastatic vs. non-metastatic, resistant vs. non-resistant) by integrating regulatory and transcriptional changes. Thus, we provide a new way to discover key molecular drivers underlying disease state transitions at single-cell resolution.

Although the proposed method demonstrates promising performance across different datasets and experimental setting, there are still some limitations that could be further improved. First, our method use prior regulatory knowledge as guidance for GRN inference. While this strategy helps improve inference, its effectiveness depends on the nature of the prior knowledge itself. Existing prior networks in public databases often have limited cell-type specificity, which may result in a supervisory signal that is not fully representative of the target cellular system. In future work, we plan to incorporate other omics information, such as chromatin accessibility. Different omics data may provide complementary perspectives on cell–type-specific regulatory relation-ships. Second, the current model primarily relies on graph neural networks to capture information propagation between genes. This allows the model to learn regulatory patterns from co-expression data and prior networks. However, our model could not distinguish direct regulations from indirect effects. Future efforts should incorporate causal inference or perturbation modeling strategies[47] to enable a more refined interpretation of direct versus indirect regulatory relationships.

In conclusion, we proposed a GRN inference model based on multi-channel graph neural network architecture. Our method effectively combines coexpression patterns with topological priors and maintains robust performance. Building upon this algorithm, we further developed a computational framework to jointly analyze regulatory structure changes and differential gene expression. The proposed method offers a new avenue for discovering key molecular drivers underlying disease state transitions.

## 5. Acknowledgements

We are grateful to the High Performance Computing Center of Central South University for partial support of this work.

## 6. Declaration of competing interest

The authors affirm that they possess no known competing interests that could potentially appear to affect the work presented in this paper.

## 7. Data availability

The source code of the proposed method is available at https://github.com/LiangXujun/DFGNN

## Notes

### Competing Interest Statement

The authors have declared no competing interest.

## References

[1] M. Smith, P. L. Flodman, Expanded insights into mechanisms of gene expression and disease related disruptions, Frontiers in Molecular Biosciences 5 (Nov. 2018). doi:10.3389/fmolb.2018.00101.

[2] L. E. Chai, S. K. Loh, S. T. Low, M. S. Mohamad, S. Deris, Z. Zakaria, A review on the computational approaches for gene regulatory network construction, Computers in Biology and Medicine 48 (2014) 55–65. doi:10.1016/j.compbiomed.2014.02.011.

[3] D. Arendt, J. M. Musser, C. V. H. Baker, A. Bergman, C. Cepko, D. H. Erwin, M. Pavlicev, G. Schlosser, S. Widder, M. D. Laubichler, G. P. Wagner, The origin and evolution of cell types, Nature Reviews Genetics 17 (12) (2016) 744–757. doi:10.1038/nrg.2016.127.

[4] M. W. E. J. Fiers, L. Minnoye, S. Aibar, C. Bravo González-Blas, Z. Kalender Atak, S. Aerts, Mapping gene regulatory networks from single-cell omics data, Briefings in Functional Genomics 17 (4) (2018) 246–254. doi:10.1093/bfgp/elx046.

[5] S. van Dam, U. Võsa, A. van der Graaf, L. Franke, J. P. de Magalhães, Gene co-expression analysis for functional classification and gene–disease predictions, Briefings in Bioinformatics (2017) bbw139.doi:10.1093/bib/bbw139.

[6] V. A. Huynh-Thu, A. Irrthum, L. Wehenkel, P. Geurts, Inferring regulatory networks from expression data using tree-based methods, PLoS ONE 5 (9) (2010) e12776. doi:10.1371/journal.pone.0012776.

[7] T. Moerman, S. Aibar Santos, C. Bravo González-Blas, J. Simm, Y. Moreau, J. Aerts, S. Aerts, Grnboost2 and arboreto: efficient and scalable inference of gene regulatory networks, Bioinformatics 35 (12) (2018) 2159–2161. doi:10.1093/bioinformatics/bty916.

[8] S. Aibar, C. B. González-Blas, T. Moerman, V. A. Huynh-Thu, H. Imrichova, G. Hulselmans, F. Rambow, J.-C. Marine, P. Geurts, J. Aerts, J. van den Oord, Z. K. Atak, J. Wouters, S. Aerts, Scenic: single-cell regulatory network inference and clustering, Nature Methods 14 (11) (2017) 1083–1086. doi:10.1038/nmeth.4463.

[9] H. Shu, J. Zhou, Q. Lian, H. Li, D. Zhao, J. Zeng, J. Ma, Modeling gene regulatory networks using neural network architectures, Nature Computational Science 1 (7) (2021) 491–501. doi:10.1038/s43588-021-00099-8.

[10] G. Mao, J. Liu, An unsupervised deep learning framework for gene regulatory network inference from single-cell expression data, in: 2023 IEEE International Conference on Bioinformatics and Biomedicine (BIBM), IEEE, 2023, pp. 2663–2670. doi:10.1109/bibm58861.2023.10385528.

[11] J. Wang, Y. Chen, Q. Zou, Inferring gene regulatory network from single-cell transcriptomes with graph autoencoder model, PLOS Genetics 19 (9) (2023) e1010942. doi:10.1371/journal.pgen.1010942.

[12] J. Chen, C. Cheong, L. Lan, X. Zhou, J. Liu, A. Lyu, W. K. Cheung, L. Zhang, Deepdrim: a deep neural network to reconstruct cell-type-specific gene regulatory network using single-cell rna-seq data, Briefings in Bioinformatics 22 (6) (Aug. 2021). doi:10.1093/bib/bbab325.

[13] Z.-H. Du, Y.-H. Wu, Y.-A. Huang, J. Chen, G.-Q. Pan, L. Hu, Z.-H. You, J.-Q. Li, Graphtgi: an attention-based graph embedding model for predicting tf-target gene interactions, Briefings in Bioinformatics 23 (3) (Apr. 2022). doi:10.1093/bib/bbac148.

[14] G. Chen, Z.-P. Liu, Graph attention network for link prediction of gene regulations from single-cell rna-sequencing data, Bioinformatics 38 (19) (2022) 4522–4529. doi:10.1093/bioinformatics/btac559.

[15] S. Wu, K. Jin, M. Tang, Y. Xia, W. Gao, Inference of gene regulatory networks based on multi-view hierarchical hypergraphs, Interdisciplinary Sciences: Computational Life Sciences 16 (2) (2024) 318–332. doi:10.1007/s12539-024-00604-3.

[16] Y. Gan, J. Yu, G. Xu, C. Yan, G. Zou, Inferring gene regulatory networks from single-cell transcriptomics based on graph embedding, Bioinformatics 40 (5) (May 2024). doi:10.1093/bioinformatics/btae291.

[17] A. Wagner, A. Regev, N. Yosef, Revealing the vectors of cellular identity with single-cell genomics, Nature Biotechnology 34 (11) (2016) 1145–1160. doi:10.1038/nbt.3711.

[18] A. Pratapa, A. P. Jalihal, J. N. Law, A. Bharadwaj, T. M. Murali, Benchmarking algorithms for gene regulatory network inference from single-cell transcriptomic data, Nature Methods 17 (2) (2020) 147–154. doi:10.1038/s41592-019-0690-6.

[19] L.-F. Chu, N. Leng, J. Zhang, Z. Hou, D. Mamott, D. T. Vereide, J. Choi, C. Kendziorski, R. Stewart, J. A. Thomson, Single-cell rna-seq reveals novel regulators of human embryonic stem cell differentiation to definitive endoderm, Genome Biology 17 (1) (Aug. 2016). doi:10.1186/s13059-016-1033-x.

[20] J. G. Camp, K. Sekine, T. Gerber, H. Loeffler-Wirth, H. Binder, M. Gac, S. Kanton, J. Kageyama, G. Damm, D. Seehofer, L. Belicova, M. Bickle, R. Barsacchi, R. Okuda, E. Yoshizawa, M. Kimura, H. Ayabe, H. Taniguchi, T. Takebe, B. Treutlein, Multilineage communication regulates human liver bud development from pluripotency, Nature 546 (7659) (2017) 533–538. doi:10.1038/nature22796.

[21] T. Hayashi, H. Ozaki, Y. Sasagawa, M. Umeda, H. Danno, I. Nikaido, Single-cell full-length total rna sequencing uncovers dynamics of recursive splicing and enhancer rnas, Nature Communications 9 (1) (Feb. 2018). doi:10.1038/s41467-018-02866-0.

[22] A. K. Shalek, R. Satija, J. Shuga, J. J. Trombetta, D. Gennert, D. Lu, P. Chen, R. S. Gertner, J. T. Gaublomme, N. Yosef, S. Schwartz, B. Fowler, S. Weaver, J. Wang, X. Wang, R. Ding, R. Raychowdhury, N. Friedman, N. Hacohen, H. Park, A. P. May, A. Regev, Single-cell rna-seq reveals dynamic paracrine control of cellular variation, Nature 510 (7505) (2014) 363–369. doi:10.1038/nature13437.

[23] S. Nestorowa, F. K. Hamey, B. Pijuan Sala, E. Diamanti, M. Shepherd, E. Laurenti, N. K. Wilson, D. G. Kent, B. Göttgens, A single-cell resolution map of mouse hematopoietic stem and progenitor cell differentiation, Blood 128 (8) (2016) e20.#x2013;e31. doi:10.1182/blood-2016-05-716480.

[24] D. Szklarczyk, A. L. Gable, D. Lyon, A. Junge, S. Wyder, J. Huerta-Cepas, M. Simonovic, N. T. Doncheva, J. H. Morris, P. Bork, L. J. Jensen, C. Mering, String 11: pprotein–protein association networks with increased coverage, supporting functional discovery in genome-wide experimental datasets, Nucleic Acids Research 47 (D1) (2018) D607–D613. doi:10.1093/nar/gky1131.

[25] L. Garcia-Alonso, C. H. Holland, M. M. Ibrahim, D. Turei, J. Saez-Rodriguez, Benchmark and integration of resources for the estimation of human transcription factor activities, Genome Research 29 (8) (2019) 1363–1375. doi:10.1101/gr.240663.118.

[26] Z.-P. Liu, C. Wu, H. Miao, H. Wu, Regnetwork: an integrated database of transcriptional and post-transcriptional regulatory networks in human and mouse, Database 2015 (2015) bav095. doi:10.1093/database/bav095.

[27] H. Han, J.-W. Cho, S. Lee, A. Yun, H. Kim, D. Bae, S. Yang, C. Y. Kim, M. Lee, E. Kim, S. Lee, B. Kang, D. Jeong, Y. Kim, H.-N. Jeon, H. Jung, S. Nam, M. Chung, J.-H. Kim, I. Lee, Trrust 2: an expanded reference database of human and mouse transcriptional regulatory interactions, Nucleic Acids Research 46 (D1) (2017) D380–D386. doi:10.1093/nar/gkx1013.

[28] J. Gasteiger, A. Bojchevski, S. Günnemann, Predict then propagate: Graph neural networks meet personalized pagerank, International Conference on Learning Representations (ICLR), New Orleans, LA, USA, 2019 (Oct. 2018). 1810.05997, doi:10.48550/ARXIV.1810.05997.

[29] X. Wang, M. Zhu, D. Bo, P. Cui, C. Shi, J. Pei, Am-gcn: Adaptive multi-channel graph convolutional networks, in: Proceedings of the 26th ACM SIGKDD International Conference on Knowledge Discovery and Data Mining, KDD ‘20, ACM, 2020, pp. 1243–1253. doi:10.1145/3394486.3403177.

[30] R. T. Davis, K. Blake, D. Ma, M. B. I. Gabra, G. Hernandez, A. T. Phung, Y. Yang, D. Maurer, A. E. Y. T. Lefebvre, H. Alshetaiwi, Z. Xiao, J. Liu, J. W. Locasale, M. A. Digman, E. Mjolsness, M. Kong, Z. Werb, D. A. Lawson, Transcriptional diversity and bioenergetic shift in human breast cancer metastasis revealed by single-cell rna sequencing, Nature Cell Biology 22 (2020) 310 – 320. URL https://api.semanticscholar.org/CorpusID:212580424

[31] Y. C. Cohen, M. Zada, S.-Y. Wang, C. Bornstein, E. David, A. Moshe, B. Li, S. Shlomi-Loubaton, M. E. Gatt, C. Gur, N. Lavi, C. Ganzel, E. Luttwak, E. Chubar, O. Rouvio, I. Vaxman, O. Pasvolsky, M. Ballan, T. Tadmor, A. Nemets, O. Jarchowcky-Dolberg, O. Shvetz, M. Laiba, O. Shpilberg, N. Dally, I. Avivi, A. Weiner, I. Amit, Identification of resistance pathways and therapeutic targets in relapsed multiple myeloma patients through single-cell sequencing, Nature Medicine 27 (3) (2021) 491–503. doi:10.1038/s41591-021-01232-w.

[32] Y. Hao, T. Stuart, M. H. Kowalski, S. Choudhary, P. Hoffman, A. Hartman, A. Srivastava, G. Molla, S. Madad, C. Fernandez-Granda, R. Satija, Dictionary learning for integrative, multimodal and scalable single-cell analysis, Nature Biotechnology 42 (2) (2023) 293–304. doi:10.1038/s41587-023-01767-y. URL https://doi.org/10.1038/s41587-023-01767-y

[33] B. Li, C. Wang, Y. Wang, P. Li, Z.-P. Liu, Regnetwork 2025: an integrative data repository for gene regulatory networks in human and mouse, Nucleic Acids Research 54 (D1) (2025) D1234–D1241. doi:10.1093/nar/gkaf779.

[34] Z. Fang, X. Liu, G. Peltz, Gseapy: a comprehensive package for performing gene set enrichment analysis in python, Bioinformatics 39 (1) (Nov. 2022). doi:10.1093/bioinformatics/btac757.

[35] B. Perozzi, R. Al-Rfou, S. Skiena, Deepwalk: online learning of social representations, in: Proceedings of the 20th ACM SIGKDD international conference on Knowledge discovery and data mining, KDD ‘14, ACM, 2014, pp. 701–710. doi:10.1145/2623330.2623732.

[36] K. Hassani, A. H. K. Ahmadi, Contrastive multi-view representation learning on graphs, in: International Conference on Machine Learning, 2020. URL https://api.semanticscholar.org/CorpusID:219558740

[37] G. Mao, Z. Pang, K. Zuo, Q. Wang, X. Pei, X. Chen, J. Liu, Predicting gene regulatory links from single-cell rna-seq data using graph neural networks, Briefings in Bioinformatics 24 (6) (Sep. 2023). doi:10.1093/bib/bbad414.

[38] B. Buyuk, S. Jin, K. Ye, Epithelial-to-mesenchymal transition signaling pathways responsible for breast cancer metastasis, Cellular and Molecular Bioengineering 15 (1) (2021) 1–13. doi:10.1007/s12195-021-00694-9.

[39] K. Milde-Langosch, The fos family of transcription factors and their role in tumourigenesis, European Journal of Cancer 41 (16) (2005) 2449–2461. doi:10.1016/j.ejca.2005.08.008.

[40] D. Zimmerli, C. S. Brambillasca, F. Talens, J. Bhin, R. Linstra, L. Romanens, A. Bhattacharya, S. E. P. Joosten, A. M. Da Silva, N. Padrao, M. D. Wellenstein, K. Kersten, M. de Boo, M. Roorda, L. Henneman, R. de Bruijn, S. Annunziato, E. van der Burg, A. P. Drenth, C. Lutz, T. Endres, M. van de Ven, M. Eilers, L. Wessels, K. E. de Visser, W. Zwart, R. S. N. Fehrmann, M. A. T. M. van Vugt, J. Jonkers, Myc promotes immune-suppression in triple-negative breast cancer via inhibition of interferon signaling, Nature Communications 13 (1) (Nov. 2022). doi:10.1038/s41467-022-34000-6.

[41] E. M. Sewify, O. A. Afifi, E. Mosad, A. H. Zaki, S. A. El Gammal, Cyclin d1 amplification in multiple myeloma is associated with multidrug resistance expression, Clinical Lymphoma Myeloma and Leukemia 14 (3) (2014) 215–222. doi:10.1016/j.clml.2013.07.008.

[42] Y. Yuan, M. Guo, C. Gu, Y. Yang, The role of wnt/beta-catenin signaling pathway in the pathogenesis and treatment of multiple myeloma (review)., American journal of translational research 13 (2021) 9932–9949.

[43] M. Kobune, H. Chiba, J. Kato, K. Kato, K. Nakamura, Y. Kawano, K. Takada, R. Takimoto, T. Takayama, H. Hamada, Y. Niitsu, Wnt3/rhoa/rock signaling pathway is involved in adhesion-mediated drug resistance of multiple myeloma in an autocrine mechanism., Molecular cancer therapeutics 6 (2007) 1774–1784. doi:10.1158/1535-7163.MCT-06-0684.

[44] O. Faruq, D. Zhao, M. Shrestha, A. Vecchione, E. Zacksenhaus, H. Chang, Targeting an mdm2/myc axis to overcome drug resistance in multiple myeloma., Cancers 14 (Mar. 2022). doi:10.3390/cancers14061592.

[45] T. A. Bloedjes, G. de Wilde, G. H. Khan, T. C. Ashby, J. D. Shaughnessy, F. Zhan, R. H. Houtkooper, R. J. Bende, C. J. M. van Noesel, M. Spaargaren, J. E. J. Guikema, Akt supports the metabolic fitness of multiple myeloma cells by restricting foxo activity., Blood advances 7 (2023) 1697–1712. doi:10.1182/bloodadvances.2022007383.

[46] X. Liu, J. Yi, T. Li, J. Wen, K. Huang, J. Liu, G. Wang, P. Kim, Q. Song, X. Zhou, Drmref: comprehensive reference map of drug resistance mechanisms in human cancer., Nucleic acids research 52 (2024) D1253–D1264. doi:10.1093/nar/gkad1087.

[47] Y. Ji, M. Lotfollahi, F. A. Wolf, F. J. Theis, Machine learning for perturbational single-cell omics., Cell systems 12 (2021) 522–537. doi:10.1016/j.cels.2021.05.016.

